# DNA damage is a major cause of sequencing errors, directly confounding variant identification

**DOI:** 10.1101/070334

**Authors:** Lixin Chen, Pingfang Liu, Thomas C. Evans, Laurence M. Ettwiller

## Abstract

Pervasive mutations in somatic cells generate a heterogeneous genomic population within an organism and may result in serious medical conditions. While cancer is the most studied disease associated with somatic variations, recent advances in single cell and ultra deep sequencing indicate that a number of phenotypes and pathologies are impacted by cell specific variants. Currently, the accurate identification of low allelic frequency somatic variants relies on a combination of deep sequencing coverage and multiple evidences of the presence of variants. However, in this study we show that false positive variants can account for more than 70% of identified somatic variations, rendering conventional detection methods inadequate for accurate determination of low allelic variants. Interestingly, these false positive variants primarily originate from mutagenic DNA damage which directly confounds determination of genuine somatic mutations. Furthermore, we developed and validated a simple metric to measure mutagenic DNA damage, and demonstrated that mutagenic DNA damage is the leading cause of sequencing errors in widely used resources including the 1000 Genomes Project and The Cancer Genome Atlas.

## Introduction

Each somatic cell within an organism contains sequence variants that are either unique or shared with only a few other cells [1,2]. Somatic variants alter the germline genome one cell at a time and, in some cases, lead to pathological conditions including cancer [3]. Thus, the accurate identification of tumor-associated variants is important for the proper diagnosis and prognosis of cancer, holding the potential to direct personalized treatments. Next generation sequencing (NGS) technologies have been pivotal for the systematic identification and characterization of these variants. Nonetheless, owing to tumor heterogeneity or contamination by normal cells, somatic variants in cancer are often found at low allelic frequencies [4,5] and thus, their identification remains a challenge.

Beyond tumor detection, a number of applications rely on the accurate detection of low allelic variants typically found in heterogeneous samples. Detection of such variants is achieved by deep sequencing of the sample. Artifactual variants resulting from sequencing errors are confounding the identification of real variants of low allelic frequency. The prevalence of these artifactual errors defines the threshold level of low allelic variant detection. The majority of sequencing errors are believed to be caused by polymerases incorporating an incorrect nucleotide base during amplification or incorrect base calling during sequencing [6]. Thus, significant time and energy has been invested to improve polymerase fidelity and sequencing accuracy. Meanwhile, mutagenic DNA damage has been recognized as a major source of sequencing error in specialized samples such as FFPE DNA [7], ancient DNA [8] and more recently circulating tumor DNA [9]. Another study has shown that a common technique used in sample preparation for DNA sequencing induces oxidative damage [10] raising the possibility that sequencing high quality human genomic DNA may also be affected by mutagenic damage.

Herein, we report that a significant number of sequencing artifacts are caused by DNA damage, and are found in essentially all genomic samples analyzed. Furthermore, the damage spectrum correlates with the procedures used for DNA storage and handling during library preparation, and confounds determination of the actual mutation spectrum found in cancers. A detailed analysis of two commonly used population resources, the 1000 Genomes Project and The Cancer Genome Atlas (TCGA) project, revealed that reported sequencing reads have an excess of G to T transversions, a signature of the oxidation of deoxyguanosine to 7,8-dihydro-8-oxoguanine (8-oxo-dG) [11], as well as further spurious errors due to damage. More importantly, we estimated that the majority of G to T transversions found in sequencing reads are due to damage for 73% of the TCGA samples. Consequently, damage alters somatic variant calling with an estimated seven wrongly annotated nonsense somatic variants in cancer genes per sample.

## Results

### The Global Imbalance Value (GIV) as a measure of DNA damage

To estimate the number of erroneous variants arising from DNA damage, we take advantage of the directional adaptors used in the Illumina library preparation workflow (*Figure 1A and Supplementary text 1*). The adaptor directionality permits paired-end sequencing, a process in which both ends of a target sequence are read independently leading to the sequencing of the template strand in the first read (R1) and the reverse complement of the template strand in the second read (R2). We further utilize the fact that a damaged base will cause an erroneous base change in only one strand of a DNA duplex. Thus, damage leading to a systematic misincorporation of a defined base opposite a damaged base (e.g. dA opposite 8-oxo-dG, which results in a G to T miscall) results in a global excess of a variant in R1 when compared to R2. This imbalance is evident when the total number of a variant in R1 is compared to the total number of the same variant type in R2 (*Figure 1A*). Based on this imbalance, we have devised an analysis strategy to deconvolute both the origin and orientation of variants and computed a novel metric, *the Global Imbalance Value* (GIV) score indicative of damage (*Supplementary text 1 and source code available at https://github.com/Ettwiller/Damageestimator*). Sequencing samples have 12 GIV scores, one per variant type. We define samples showing a GIV score above 1.5 as severely damaged. At this GIV score, there are 1.5 times more variants on R1 sequences compared to R2 sequences suggesting that at least one third of the variants are erroneous. Undamaged DNA samples have a GIV score of 1.

**Figure 1.**
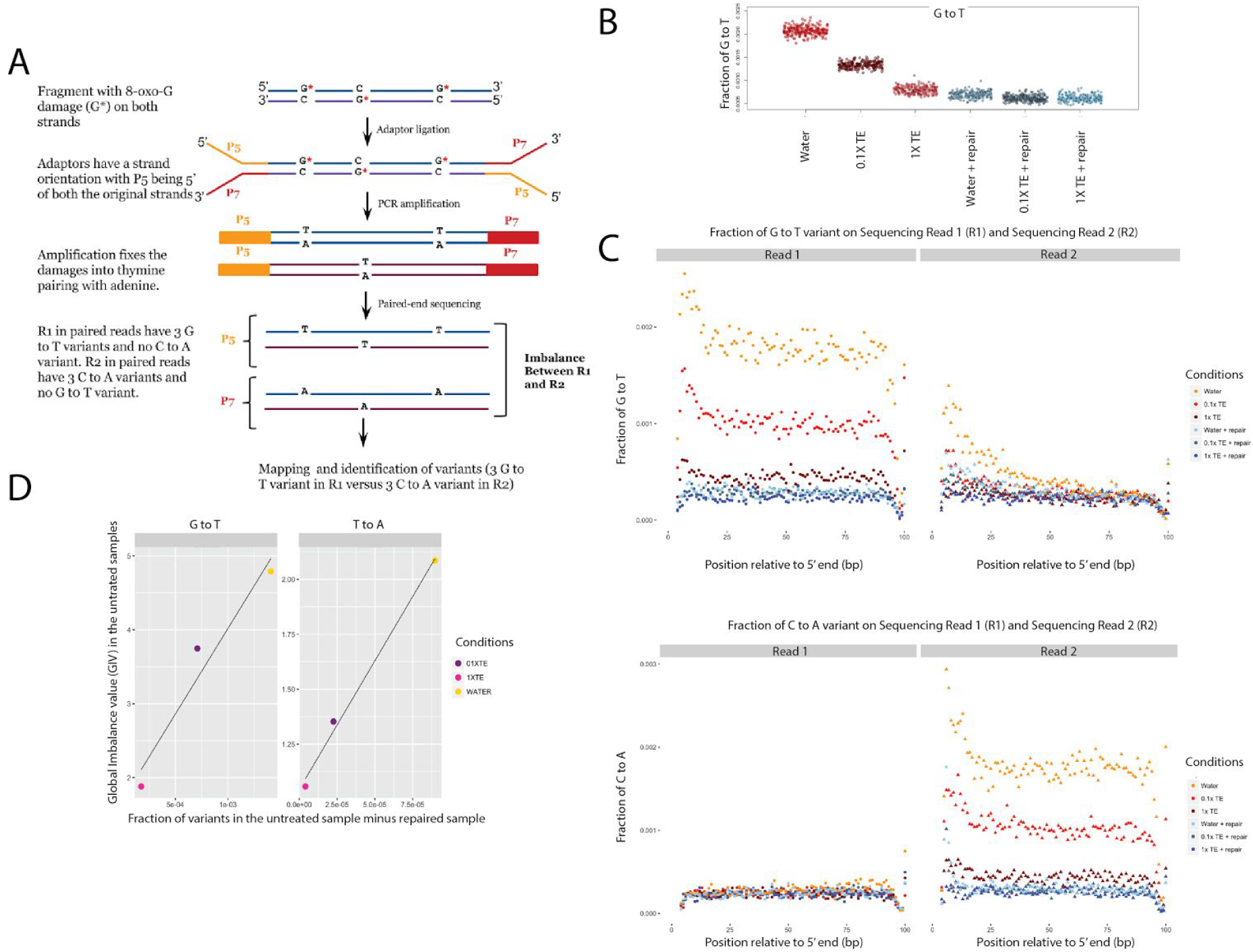
The Global Imbalance Value (GIV) score. **A.** <u>Principle of the GIV score</u>: Illumina adaptors are directional in nature, enabling consistent paired end sequencing within clusters. This property results in sequencing of the original strand orientation in the R1 reads (from the P5 adaptor) whereas the reverse complement orientation is read in the R2 reads (from the P7 adaptor). As damage affects only one base of a pair, damage such as 8-oxo-dG leads to an excess of G to T transversion errors when R1 is mapped to a reference genome, whereas, the R2 reads will show an excess of the reverse complement of G to T, i.e. C to A transversion errors, instead. As a consequence, there is a global imbalance in the number of G to T variants in R1 compared to R2 sequences. This imbalance is specific to damage and is the basis of the GIV score (see Supplementary text 1). **B.** Overall fraction of G to T variants (normalized to the total number of G) for R1 and the reverse complement of R2 sequences. Different buffers were used during acoustic shearing (x-axis). Data in red were from samples that were not repaired and data in blue were from samples that were repaired. Each point corresponds to a random sampling of 2 million sequence positions. All samples are derived from the same human genomic DNA. **C.** Variant profile: The fraction of G to T and C to A variants in R1 and R2 sequences are plotted as a function of the read (R1 or R2) and the positions on the read (in bp). **D.** Correlation (R=0.97) between the degree of damage that is repaired by the DNA repair enzyme cocktail and GIV for G to T variant (GIV_G_T_) and T to A variant (GIV_T_A_).

To experimentally validate the GIV score and provide an independent quantification of damage, we designed a pilot experiment using human genomic DNA containing various amounts of 8-oxo-dG, an oxidative damage introduced during acoustic shearing [10]. This damaged base is known to pair with adenine resulting in a G to T transversions after amplification [12]. We treated the damaged DNA with an enzyme cocktail that eliminates and repairs DNA damage prior to library preparation (*Supplementary text 2 **and Supplementary Materials and Methods***) [11]. Sequencing the same sample with and without treatment with the DNA repair enzyme cocktail quantifies the rate of erroneous variants that are specifically introduced by damage. Libraries from treated and untreated samples were paired-end sequenced using an Illumina MiSeq platform, resulting in an average of ∼4 million paired-end reads per sample. Reads were mapped to a reference human genome (hg19) using BWA-MEM, and mapped sequences were analyzed using our novel orientation-aware variant calling algorithm and GIV scoring (***Materials and Methods***).

We found that the frequency of G to T transversions in the same DNA sample varies according to the shearing conditions used and is inversely correlated with the buffer concentration used to shear the DNA (*Figure 1B and Supplementary text 3*). Treatment of the DNA sample with the repair enzyme cocktail post acoustic shearing reduced the number of G to T variants to baseline levels at all shearing conditions (*Figure 1B*). This is consistent with 8-oxo-dG damage being the cause of the excess of G to T variants in the unrepaired samples. Additional excess of G to T variants is observed within the first ∼20 bp of the sequencing read, consistent with increased oxidative damage on single stranded DNA (*Figure 1C*). Contrary to a previous study reporting a strong preference for oxidation of G in a CGG context [10], we do not observe nucleotide context specificity (*Supplementary Figure 1A*). We confirm previous findings [10] that buffer composition during shearing is the major modulator of 8-oxo-dG damage (*Supplementary Figure 2* and *Supplementary text 3*). However, we found no effect of EDTA on the levels of DNA damage. In addition, we detected several previously uncharacterized signatures of damage that correlate with buffering conditions during shearing (*Supplementary text 3 and Supplementary Figure 1*). Surprisingly, the fraction of errors due to damage increased with increasing Phred quality score cutoff (*Supplementary Figure 3A*), indicating that errors induced by damage cannot simply be removed by elevating the stringency of base calling.

**Figure 2.**
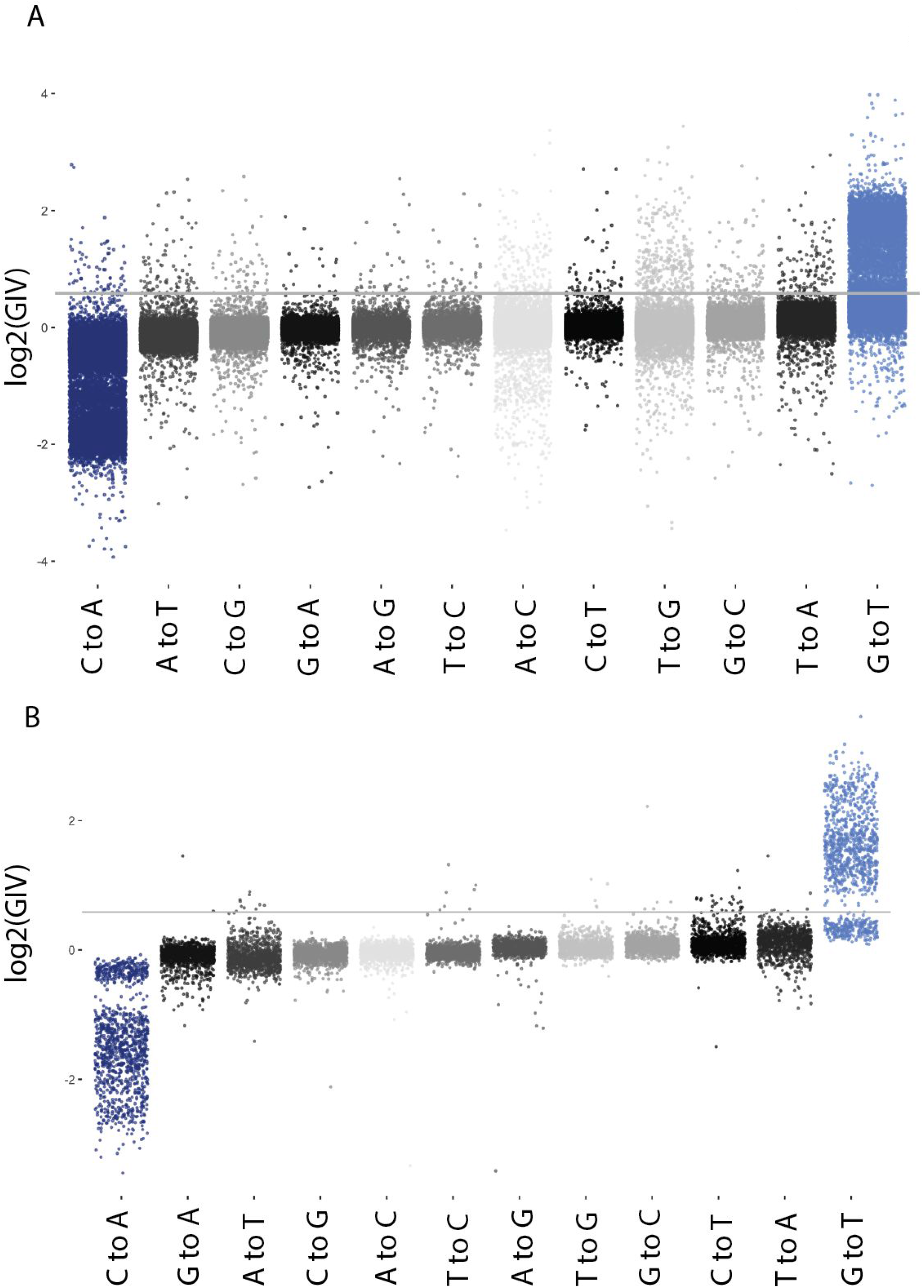
GIV scores for the 12 nucleotide substitution classes for **A.** The 1000 Genomes Project dataset and **B.** A subset of the TCGA datasets. Each point represents a single sequencing run downsampled to 5 million reads. The solid line denotes a GIV of 1.5.

**Figure 3.**
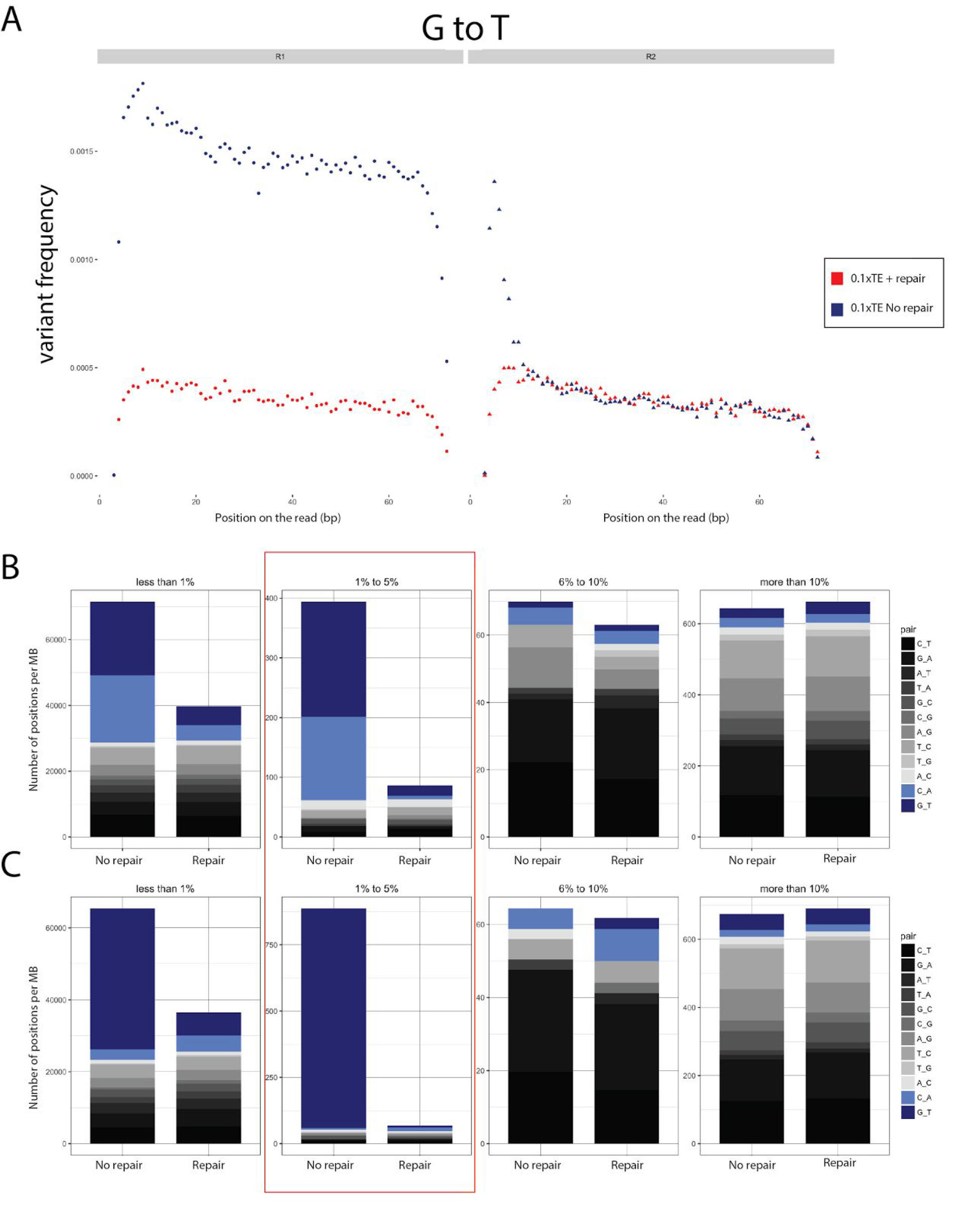
Target enrichment experiment. Genomic DNA was sheared in 0.1xTE buffer and subjected to library preparation and cancer panel target enrichment. DNA repair treatment was performed postshearing and prior to library preparation. The control represents untreated DNA. **A.** Overall G to T variant profiles across reads R1 and R2 with (red) and without (blue) repair treatment. **B.** Variant spectrum at a frequency less than 1% (left panel), 1 to 5% (center left panel), 6 to 10% (center right panel) and more than 10% (right panel). **C.** Same as for B except that only R1 reads were used for variant calling.

Importantly, the excess of G to T variants can only be observed in R1 sequences while C to A variants are found in excess in R2 sequences leading to an imbalance in variant rate and a GIV_G_T_ score>1 (*Figure 1C*). Treatment with the repair enzyme cocktail abolishes this imbalance and reduces the GIV_G_T_ score to 1. Furthermore, the GIV score correlates with the excess of the variant measured experimentally (Figure 1D), demonstrating that the GIV score can be used to accurately estimate the extent of damage in publically available datasets derived from samples that are typically not paired with undamaged controls. We estimated that the GIV score calculation is accurate at greater than 2 million reads (*Supplementary Figure 3B*).

### Damage is the major cause of sequencing error in publically available datasets

To estimate the extent of damage in public datasets, we analyzed individual sequencing runs from both the 1000 Genomes Project [13] and a subset of the TCGA dataset. For each sequencing run the GIV score was computed (**Materials and Methods**) and to our surprise both the TCGA dataset and the 1000 Genomes Project dataset showed widespread damage, particularly those leading to an excess of G to T variants (Figure 2).

Specifically, 41% of the 1000 Genomes Project datasets had a GIV_G_T_ score of at least 1.5 indicative of severely damaged samples (Figure 2A). For 73% of the TCGA datasets, the DNA showed such an extensive damage that the majority of G to T transversions are erroneous (GIV_G_T_>2), establishing damage as the leading cause of errors (Figure 2B).

The profile of damage signatures in both the 1000 Genomes Project and the TCGA datasets were similar to those obtained in our pilot experiment in which oxidative damage was introduced during library preparation. Likewise, we did not find nucleotide context specificity of G to T imbalances in the publicly available datasets (Supplementary Figure 4 A, B and C).

**Figure 4.**
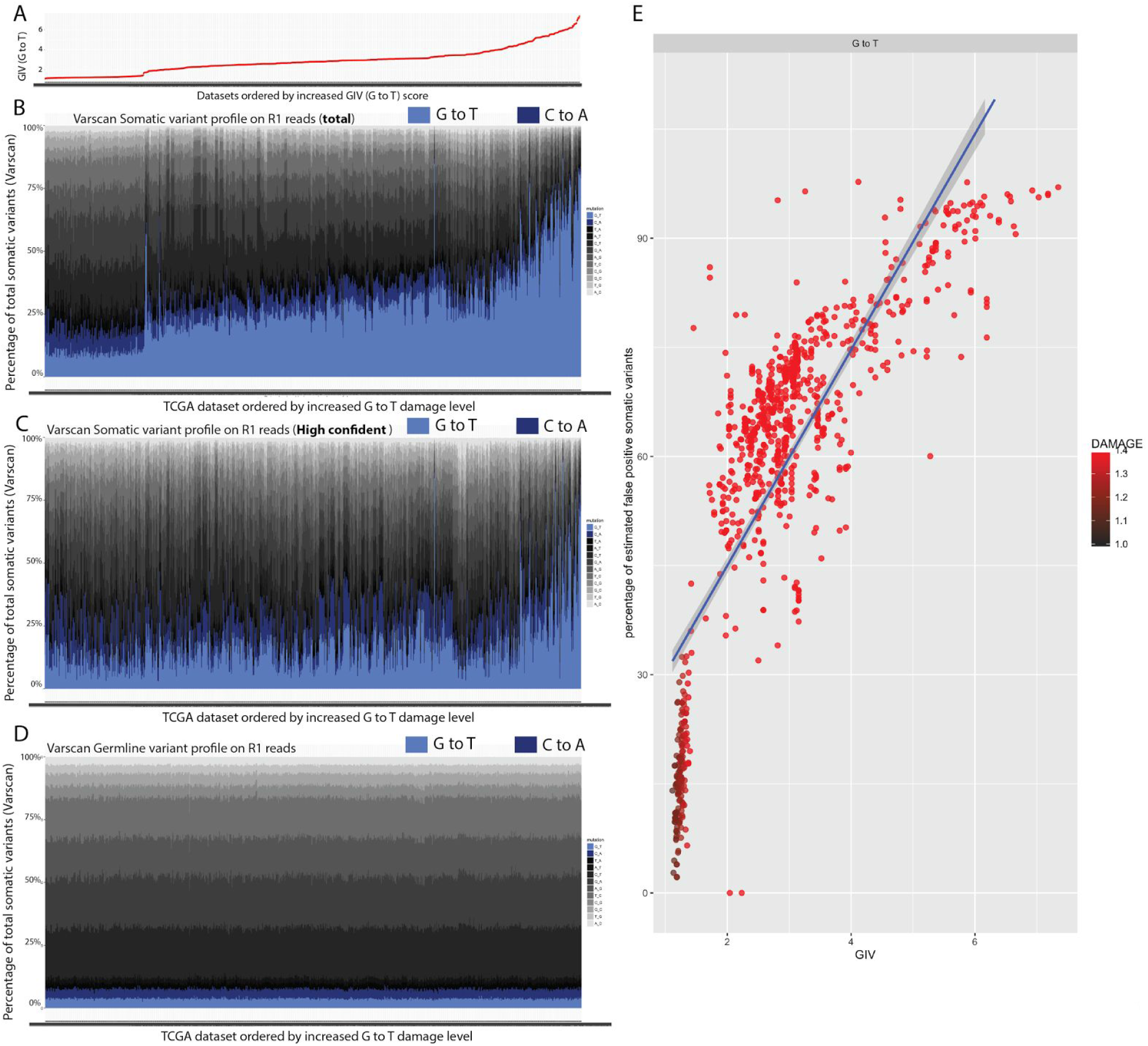
Variants identified in TGCA datasets. **A.**GIV_G_T_ scores of the ∼1800 tumor samples sorted by increasing GIV_G_T_ scores. **B.** Profiles of somatic variant calls for the ∼1800 tumor samples sorted by increasing GIV_G_T_ scores. The fraction of somatic variant calls of G to T (cyan) is higher when compared to C to A (blue) for most of the datasets and the overall fraction of somatic variant calls of G to T increases with the increase of the GIV_G_T_ score. **C.** Same as in B using the profile of high confidence somatic variant calls. **D.** Same as in B except that the profile of germline variant calls are represented. **E.** Estimated false positive rate of somatic variant calls of G to T (in percent) as a function of the GIV_G_T_ score.

Severely damaged samples leading to a T to A imbalance were detected in 0.5 % of the TCGA dataset. This T to A imbalance was also identified in our pilot experiment *(Supplemental text 3, Supplemental Figure 1B*) and similarly, the T to A variant profile found in the TCGA datasets shows context specificity and elevated variant frequency for the first ∼20 bp at the 5’ end of the R2 reads. This suggests that the damage that causes the T to A imbalance is preferentially located at the 3’ end of the original DNA fragment (Supplementary Figure 5). Other imbalances were found, including a C to T imbalance (3% of datasets), indicative of cytosine deamination. We repeated the same analysis on recent submissions to TCGA (Nov-Dec. 2015) and found similar G to T imbalances and accentuated T to A imbalances (Supplementary Figure 6).

Thus, the overwhelming majority of the publicly available dataset, including recently submitted TCGA datasets, have signatures of damage leading to erroneous calls in at least one third of the G to T variants. This result is not surprising given that the most commonly used DNA shearing approach uses buffer conditions estimated to cause a majority of damage induced G to T variants. It is therefore reasonable to assume that damage is the leading cause of sequencing errors in NGS.

### Damage leads to the incorrect identification of somatic variants

The GIV score estimates the extent of spurious variants due to damage directly on mapped reads. We know from our data that oxidative damage is a stochastic event happening at random positions with no apparent sequence context (*Supplementary Figure 1A)*. Such stochasticity implies that errors derived from damage are expected to be present at low allelic fractions. Thus we hypothesized that the identification of low frequency variants such as somatic variants is affected by damage while germline variants should remain unaffected.

To evaluate how damage affects our ability to identify somatic variants, we repeated the oxidative damage experiments using standard library preparation procedures. We further performed target-enrichment using a commercial cancer panel probe set covering about 0.79 Mb of 151 annotated cancer genes to achieve high sequencing depth at specific genomic locations (***Supplementary Materials and Methods***).

The overall measurement of damage in the enriched DNA without DNA repair is in agreement with whole genome sequencing but with slightly higher levels of damage-induced variants in the enriched DNA (GIV_G_T capture_ = 4.4 and GIV_G_T genome_ = 3.7). DNA repair fully abolished detectable damage (Figure 3A) demonstrating that the enrichment step does not drastically modify the oxidative damage rate nor introduce additional oxidative damage that would lead to G to T errors.

Next, we normalized the coverage at each genomic position to 300-fold coverage and classified variants according to frequency with very low (<1%), low to moderate (1%-5%), medium (6-10%) and high (>10%) frequency variant classes. We found that damaged DNA has an effect on very low and low to moderate frequency G to T and C to A variant positions compared to repaired DNA (Figure 3B and Supplementary table 2). More specifically, we found that DNA repair eliminates 77% and 82% of G to T and C to A variant positions in the very low and low to moderate frequency variant classes, respectively. Interestingly, when investigating the total variants profiles, the major effect of damage is reflected in the low to moderate frequency variant class. Indeed, the repaired sample shows 78% less total variants compared to the damaged sample in the moderate frequency variant class as opposed to only 44% less total variants in the very low frequency variant class (Figure 3B). We interpret this result as a high background of erroneous variants due to sequencing error and polymerase error at very low frequency (<1%), while most of the erroneous positions with low to moderate frequency variants are due to damage.

This is an important point because those genomic positions with low to moderate frequency variants are positions typically classified as somatic variants for the reason that they harbor multiple evidences of variant reads at high coverage. Looking only at the 0.79 Mb region included in the cancer panel, we found 195 genomic locations with low to moderate G to T variants in the unrepaired dataset and 12 in the repaired dataset. Moreover, amongst the 195 positions found in the unrepaired dataset, 50 are marked as deleterious and 7 are annotated as nonsense according to PredictSNP2 [14]. These results strongly indicate that more than 180 positions are false positives and are directly confounding the identification of real somatic variants in cancer genes. This figure corresponds to an average of around one erroneous call per cancer gene with some erroneous changes likely leading to diagnostic errors.

Focusing on the R1 sequences only, we see a strong imbalance of G to T positions compared to C to A positions for both the low and low to moderate frequency classes in the unrepaired dataset (Figure 3C) confirming the role of damage in erroneous variant calling. As expected, high frequency variants such as SNP’s (>10%) are not affected by damage because the rate of variant calling and variant profiles are similar between the matched repaired and unrepaired samples (Figure 3B).

In summary, our data demonstrates a direct link between damage and the ability to accurately call very low and low to moderate frequency variants. With an estimated 78% false positive rate in variants from the 1 to 5% frequency range (a frequency range expected in heterogeneous tumor samples), oxidative damage introduced into high quality DNA during standard library preparation is predicted to be the major cause of erroneous identification of somatic variants.

### DNA damage observed in the TCGA dataset directly affects variant calling

To better assess the extent that damage affects somatic variant calls in actual cancer samples, we identified germline and somatic variants for all the TCGA tumor samples with matched tumor-normal pairs (see ***Supplementary Materials and Methods***). For this, we used Varscan, a software tool previously shown to detect germline and somatic variants with high sensitivity and specificity [15]. Prior to variant calling, the mapped reads were grouped into R1 and R2 reads. This distinction allowed us to assess whether the total number of somatic mutation calls were globally balanced between the two groups. Analogous to GIV, an excess of somatic mutation calls in one group compared to the other represents erroneous calls caused by damage.

We found a large excess of G to T somatic variants when compared to the rate of C to A somatic variants for most of the datasets (Figure 4A and B). Moreover, the fraction of G to T variants compared to other variants increased with the estimated damage measured by the GIV_G_T_ score (Figure 4B). Interestingly, severely damaged samples also showed an excess of high confidence G to T somatic variants demonstrating that damage affects high confidence somatic mutation calls in these samples (Figure 4C). In contrast we found that the fraction of G to T germline variants is constant across samples and showed no excess in the R1 reads (Figure 4D) as expected for high frequency variants.

Next, we estimated the false positive rate of somatic variant calls and found that 78% of tumor samples have more than 50% false positive G to T somatic variant calls. Furthermore, the percentage of false positives strongly correlated (r=0.79) with the estimated damage in tumor samples (Figure 4E). This strong correlation between damage and false positive somatic variants indicated that damage is the direct cause of erroneous identification of somatic variants.

Unexpectedly, we also identified a smaller subset of the TCGA dataset with a large excess of both total and high confidence somatic variant calls of the C to T type (Supplementary Figure 7). Taken together, these results highlight a major confounding effect of damage on somatic mutations including high confidence somatic mutation calls in the TCGA datasets.

## Conclusion

Low frequency variants are difficult to identify because stochastic sequencing errors such as errors induced by DNA damage, as described in this report, confound their identification. To distinguish actual somatic variants from artifactual variants standard strategies have been used to increase sequencing coverage, set stringent variant frequency thresholds and apply various post-processing filters in variant calling algorithms. The application of stringent filters is not a complete solution for the detection of low frequency variants because they are likely leading to false negatives. Thus, these filtering steps are inferior substitutes to increased accuracy in sequencing. This study demonstrates that certain DNA handling procedures avoid mutagenic damage and increase sequencing accuracy. Such procedures include the use of appropriate buffer conditions when preparing libraries and the use of repair enzymes to eliminate damage. It is also important to complement careful DNA manipulation with proper computational analysis tools to identify samples containing DNA damage and eliminate sequencing data when it has the potential to lead incorrect somatic variants calls and therefore incorrect diagnostic conclusions.

## Materials and Methods

### DNA sample handling and library preparation

To introduce oxidative damage, purified human liver genomic DNA samples (BioChain Institute, Inc.) were fragmented by shearing in 1x TE (10 mM Tris pH 8, 1 mM EDTA), 0.1x TE (1 mM Tris pH 8, 0.1 mM EDTA) or water using a Covaris S2 to produce 200 bp average fragment size as determined by Bioanalyzer (Agilent Technologies) analysis. For sequencing library preparation, 800 ng of fragmented human liver DNA was subjected to library preparation using the NEBNext DNA Library Prep Master Mix Set for Illumina (NEB, Inc.) with and without DNA repair using the NEBNext FFPE DNA Repair Mix and protocol (***Supplementary Materials and Methods***). For target enrichment, 250 ng of fragmented human liver DNA was subjected to library preparation using the NEBNext Ultra II DNA Library Prep Kit for Illumina (NEB, Inc) protocol with and without DNA repair. 750 ng libraries were used for target capture using the Agilent XT ClearSeq Comprehensive Cancer Panel (Agilent Technologies, ***Supplementary Materials and Methods***).

### DNA sequencing

Libraries were paired-end sequenced using the Illumina MiSeq platform with either the 75 bp or 150 bp paired-end read mode. Reads were trimmed to remove adaptor sequences and paired-end mapped to the reference human genome using BWA-MEM algorithm. In some cases, local realignment was performed using the GenomeAnalysisTKLite from GATK (IndelRealigner).

### Public datasets

#### The Cancer Genome Atlas (TCGA)

The manisfest files were downloaded using the cghub browser (https://browser.cghub.ucsc.edu/) to download the manifest file with the following apply filters: library_strategy=WXS&refassem_short_name= HG19_Broad_variant&disease_abbr=LUAD&study=phs000178’’ -o pilot2.xml -c https:/cghub.ucsc.edu/software/downloads/cghub_public.key

For the newest TCGA dataset, the manifest file was downloaded with the following filters: published_date=[NOW-6MONTHS TO NOW]&library_strategy=WXS’’ -o newest.xml -c https:/cghub.ucsc.edu/software/downloads/cghub_public.key

Resulting bam files were downsampled to 5 millions reads.

#### The 1000 Genomes Project

Only Illumina paired-end read datasets were used. FASTQ files were downloaded and downsampled to 4 million reads.

### GIV score

Paired-end reads were obtained from each sample and adaptors were trimmed using Trim-galore. Reads were paired-end mapped to the hg19 version of the human genome using BWA-MEM with default parameters (k=1). In some cases, local realignments were performed using the GenomeAnalysisTKLite from GATK (IndelRealigner).

After mapping, the map files was organized into two groups, one generated from R1 reads and the other group generated from R2 reads. Mpileup was run independently on both groups with default parameters (and -O -s -q 10 -Q 0).

The analysis of damage was done using damage_estimator (https://github.com/Ettwiller/Damageestimator). For the GIV score, Estimate_damage_location.pl was used (available at https://github.com/Ettwiller/Damageestimator/) and run using the following parameters qualityscore 30--min_coverage_limit 1.

To correlate the GIV-score with Phred score (*Supplementary Figure 3A*), the same analysis was done as described above with the parameter--qualityscore ranging from 0 to 40 (1 increment).

In the analysis of the cancer panel enrichment experiments, the--max_coverage_limit parameter was set to 5000.

## Acknowledgements

We would like to thank Sharon Kaiser, Aaron Messelaar, Tamas Vincze and Ching Lin for IT support and compliance with the TCGA hosting guideline; Laurie Mazzola, Joanna Bybee and Danielle Rivizzigno for sequencing. We would also like to thank Richard Roberts, William Jack, Tilde Carlow, Andrew Gardner and Salvatore Russello for critical comments of the manuscript. The results shown here are in part based upon data generated by the 1000 Genomes Projects and the TCGA Research Network: http://cancergenome.nih.gov/.”

## Supplementary Materials

### Supplementary text 1. The Global Imbalance Value (GIV)

In Illumina sequencing, the P5 flow cell binding site adaptors are ligated to the 5’ end of the original DNA fragment while the P7 adaptors are ligated to the 3’ end of the original DNA fragment. Amplification during PCR or clustering preserves this orientation.

After paired-end sequencing, R1 reads always provides the sequence of the original template molecule and R2 reads always provides the sequence of the reverse complement of original template molecule. For example, 8-oxo-dG is a well-known result of oxidative damage often misread by a polymerase as a thymine (T) instead of a guanine (G). Consequently the polymerase incorporates an adenine (A) to pair with 8-oxo-dG. The erroneous variant of 8-oxo-dG is therefore captured as a G to T on R1 reads, while R2 reads capture the reverse complement variant, C to A.

Large difference in the number of G to T variants originating from the R1 reads compared to the number of G to T variants from the R2 reads reflects an imbalance, which is characteristic of damage. Mirroring this imbalance, the C to A variants show the inverse trend when compared to G to T variants (thus, the GIV score could be extended to single-end read sequencing by comparing the rate of G to T variants with the rate of C to A variants originating from the single-end reads).

We can combine these differences and calculate the global imbalance value (GIV) score using the following equation:

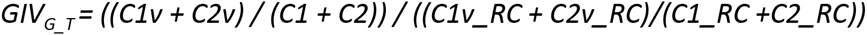

With *C1v* = Number of G to T variants in R1; *C1* = Total number of G in R1; *C2v* = Number of C to A variants in R2; *C2* = Total number of C in R2; *C1v_RC* = Number of C to A variants in R1; *C1_RC* = Total number of C in R1; *C2v_RC* = Number of G to T variants in R2; *C2_RC* = Total number of G in R2

### Supplementary text 2. 8-oxo-dG is efficiently repaired by *in vitro* DNA Repair

We used the NEBNext FFPE DNA Repair Mix (New England Biolabs, Inc.) protocol for *in vitro* DNA Repair. The general strategy for DNA repair by this mix and the related PreCR mix involves two steps: First a DNA glycosylase recognizes a specific damage and cleaves the N-glycosidic bond between the damaged base and the sugar phosphate backbone of the DNA. There are four different glycosidases, each one recognizing a particular type of DNA damages. For 8-oxo-dG damage, the glycosylase used is formamidopyrimidine [fapy]-DNA glycosylase (FPG) [11]. The cleavage by the specific glycosidase generates an apyrimidinic/apurinic (AP) site in the DNA. Alternatively, AP sites can also arise by spontaneous hydrolysis of the N-glycosidic bond. In either case, the AP site is subsequently processed by *E. coli* Endonuclease IV, which cleaves the phosphodiester backbone immediately 5’ to the AP site, resulting in a 3’ hydroxyl group and a transient 5’ abasic deoxyribose phosphate (dRP). The second step consists of the removal of the dRP, accomplished by the action of the DNA polymerase, which adds nucleotides to the 3’ end of the nick and removes the dRP moiety via its associated flap endonuclease activity. Finally the nick produced is sealed by Taq DNA ligase, thus restoring the integrity of the DNA.

### Supplementary text 3. Shearing conditions and oxidative damage

To identify the causative agent for 8-oxo-dG damage during library preparation, we subjected human genomic DNA to various shearing conditions (Supplementary table 1). Based on the degree of G to T imbalance, we found that the 10 mM Tris pH 8 +1 mM EDTA (1X TE) shearing buffer minimized oxidative damage but does not completely abolish it. We found that shearing in a 10 mM Tris, pH 8 buffer alone is sufficient to bring the G to T imbalance down to almost the same level as 1X TE, demonstrating that the buffering capacity of the solution during acoustic shearing is the major modulator of damage. The addition of EDTA in the Tris buffer marginally decreases the G to T imbalance likely due to the buffering capacity of EDTA. Industry standard for DNA shearing is in 1 mM Tris pH 8 + 0.1 mM EDTA (0.1X TE). In this buffer, the damage is estimated to be about half of the damage observed when the sample is sheared in water but still accounts for the majority of the G to T variants (Supplementary figure 2).

**Supplementary Figure 1.**
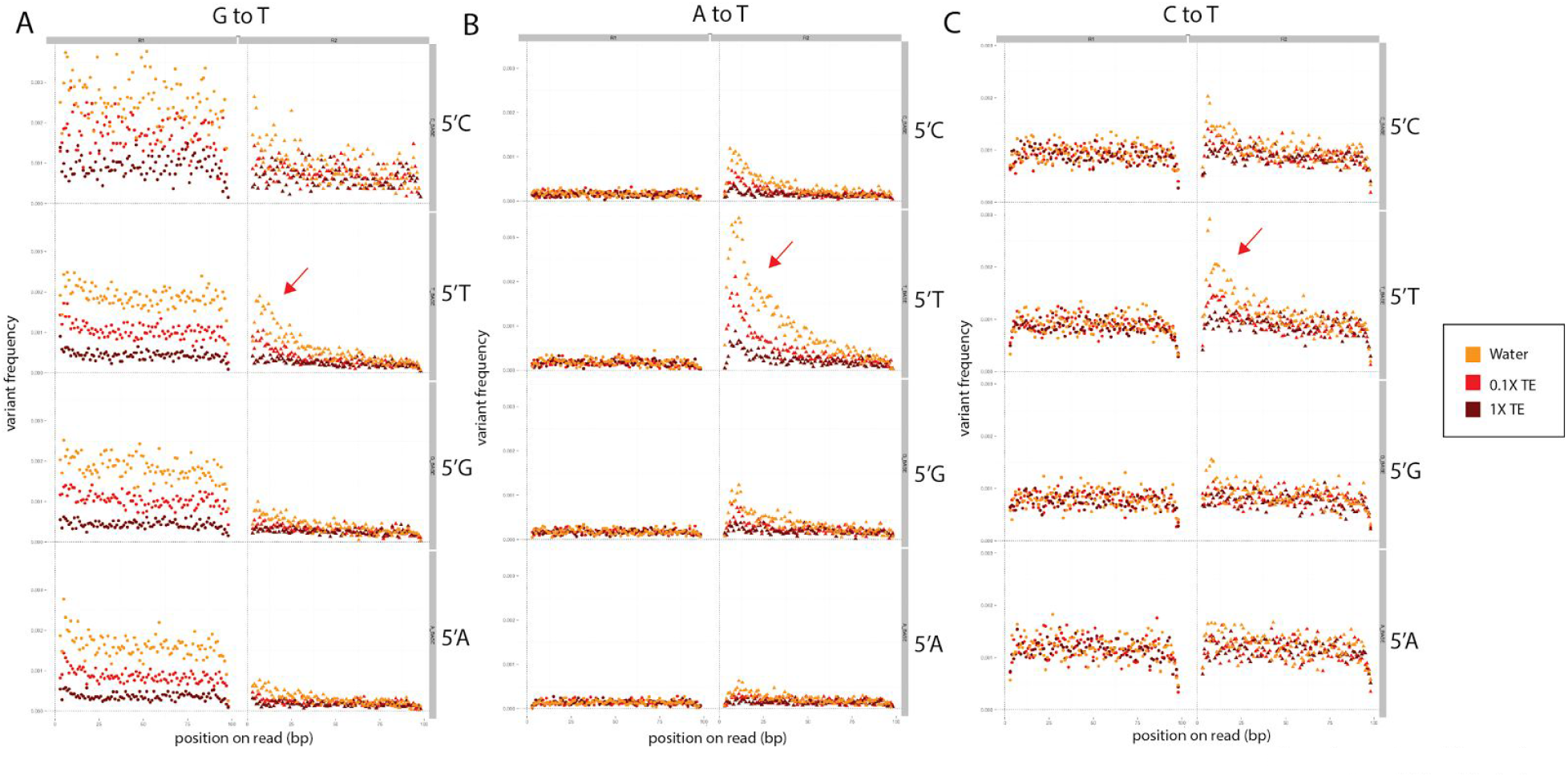
Variant profiles and context specificity across R1 and R2 sequences for G to T (**A**), A to T (**B**) and C to T (**C**) for samples sheared in water (yellow), 0.1X TE (red) and 1X TE (brown). G to T transversions are an elevated fraction of variants in R1 sequences in all contexts. In all three cases, an elevated fraction of variants (red arrow) can be found at the 5’ end of R2 sequences in a 5’ T context.

**Supplementary Figure 2.**
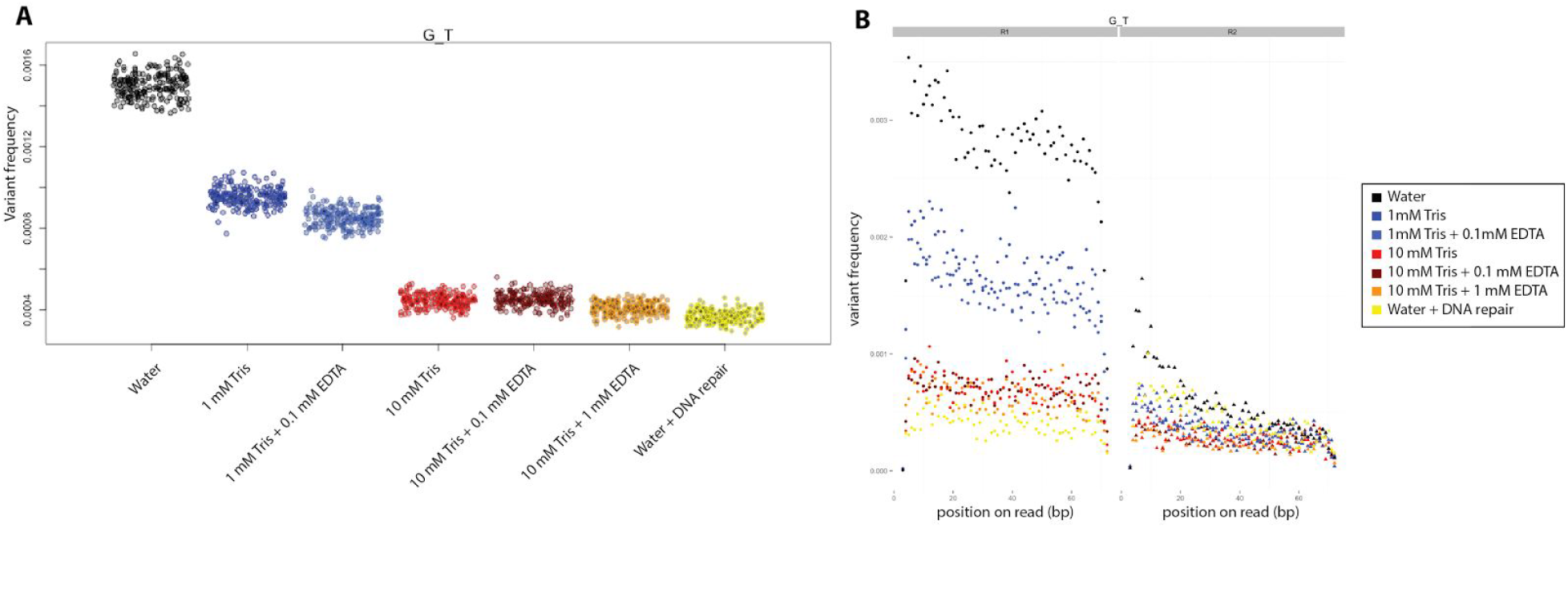
**A.** Frequency of G to T transversions in samples sheared in water (black), 1 mM Tris (dark blue), 1 mM Tris+0.1 mM EDTA (light blue), 10 mM Tris (red), 10 mM Tris+0.1 mM EDTA (dark red) and 10 mM Tris+1 mM EDTA (orange). The samples sheared in water were also treated with repair enzymes to identify the baseline G to T transversions from undamaged DNA (yellow). **B.** G to T variant frequency as a function of read position for R1 sequences and R2 sequences in samples sheared in water (black), 1 mM Tris (dark blue), 1 mM Tris+0.1 mM EDTA (light blue), 10 mM Tris (red), 10 mM Tris+0.1 mM EDTA (dark red) and 10 mM Tris+1 mM EDTA (orange). The samples sheared in water were also treated with the repair enzyme cocktail to identify the baseline G to T transversion from undamaged DNA (yellow). Imbalances between R1 and R2 sequences were identified in all conditions with the exception of the repaired sample (yellow).

**Supplementary Figure 3.**
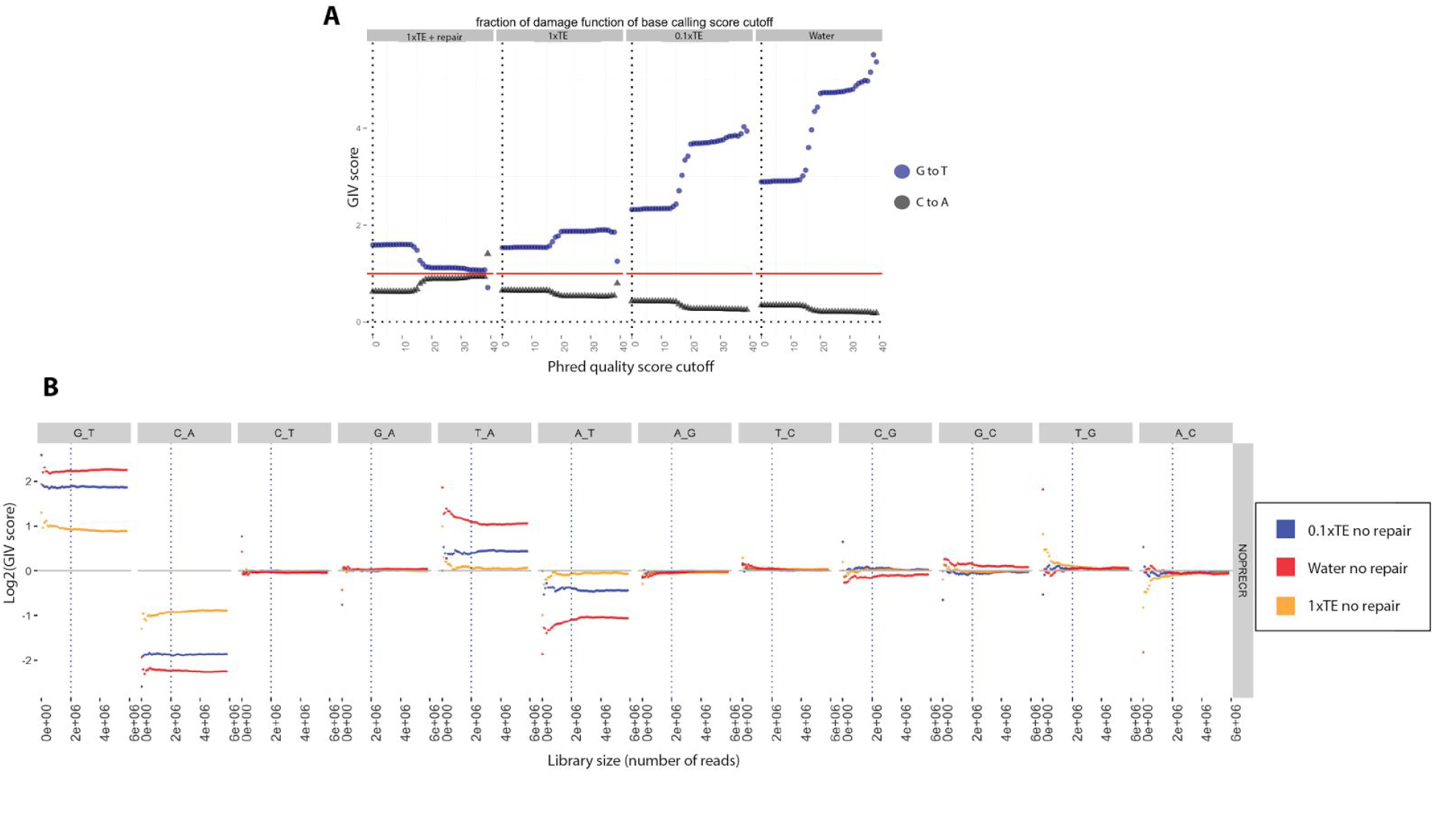
**A.** Correlation between Phred quality cutoff and GIV_G_T_ (blue) and GIV_C_A_ (black) for genomic samples sheared in various buffer conditions (Water, 0.1X TE, 1X TE) and repaired sample sheared in 1X TE. The fraction of errors due to damage increases with the confidence of base calling, explaining the increase in GIV_G_T_ with increasing phred quality cutoff in untreated samples. **B.** Effect of sequencing depth (number of reads) on GIV score. Sequencing reads from samples sonicated in water (red), 0.1X TE buffer (blue) and 1X TE buffer (orange) range from 0.1 million reads to 6 million reads. For each down-sampled experiment, GIV scores were calculated and plotted. The dotted line (at 2 million reads) indicates stabilization of the GIV score and the lower limit of reads required to accurately calculate GIV score.

**Supplementary Figure 4.**
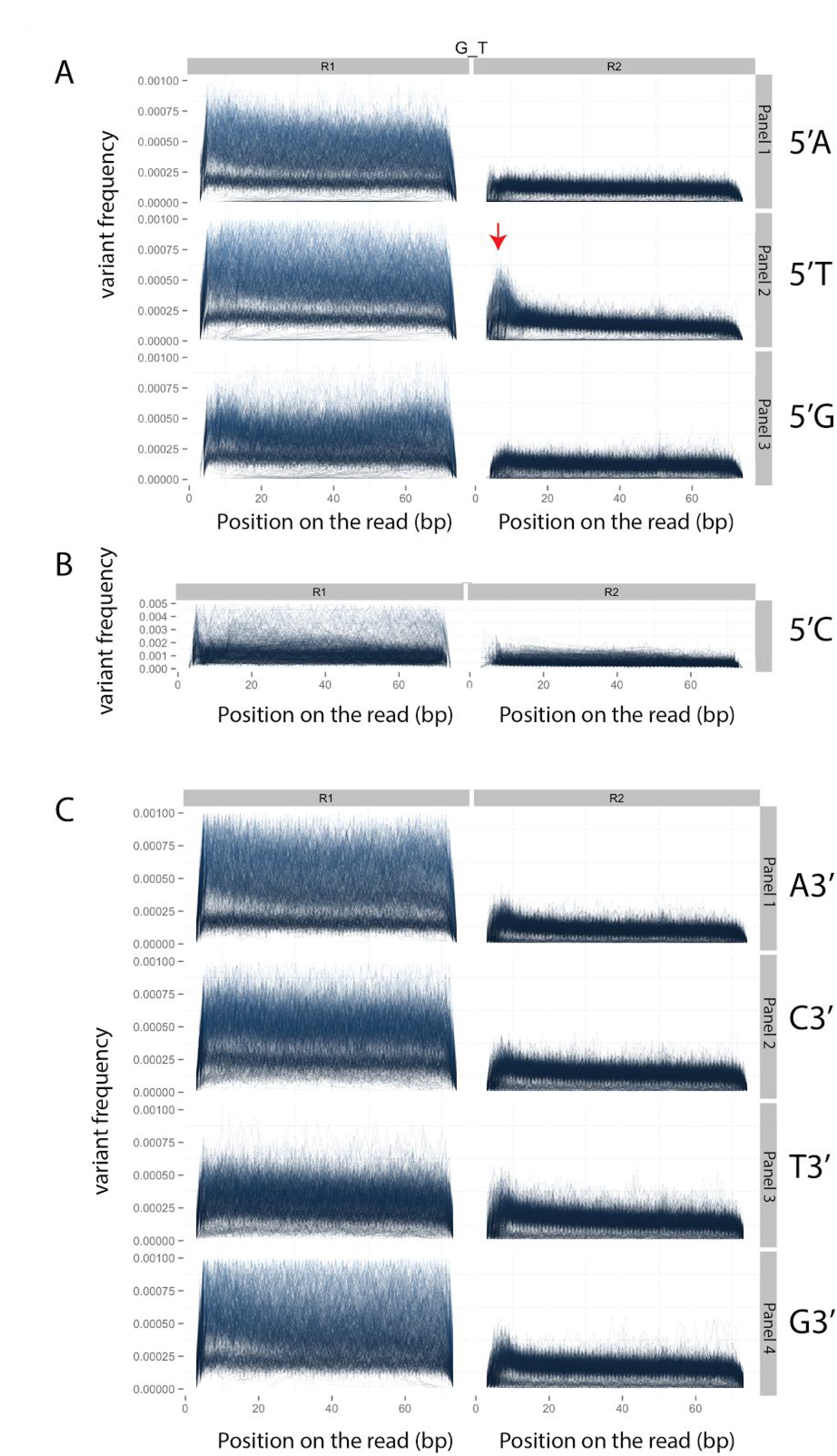
Damage in the TCGA dataset: **A.** G to T variant frequency as a function of read position in R1 and R2 for sequence context ApG (panel1), TpG (panel2) and GpG (panel3). **B.** Same as in A except for the CpG context. **C.** Same as in A except for the GpA (Panel1), GpC (panel2), GpT (panel3) and GpG (panel4) context. Each line corresponds to one TCGA sample. The imbalance of G to T on the R1 compared to R2 sequencing reads can be observed in all contexts for most of the samples. We observed an elevated G to T on the 5’ end of the R2 sequence reads corresponding to an elevated C to A on the 3’ end of the original fragments. Contrary to G to T variants of R1 sequences, this elevated C to A frequency has a strong sequence context at the TpG context (red arrow) and position specificity within the first 20 bp of the R2 reads.

**Supplementary Figure 5.**
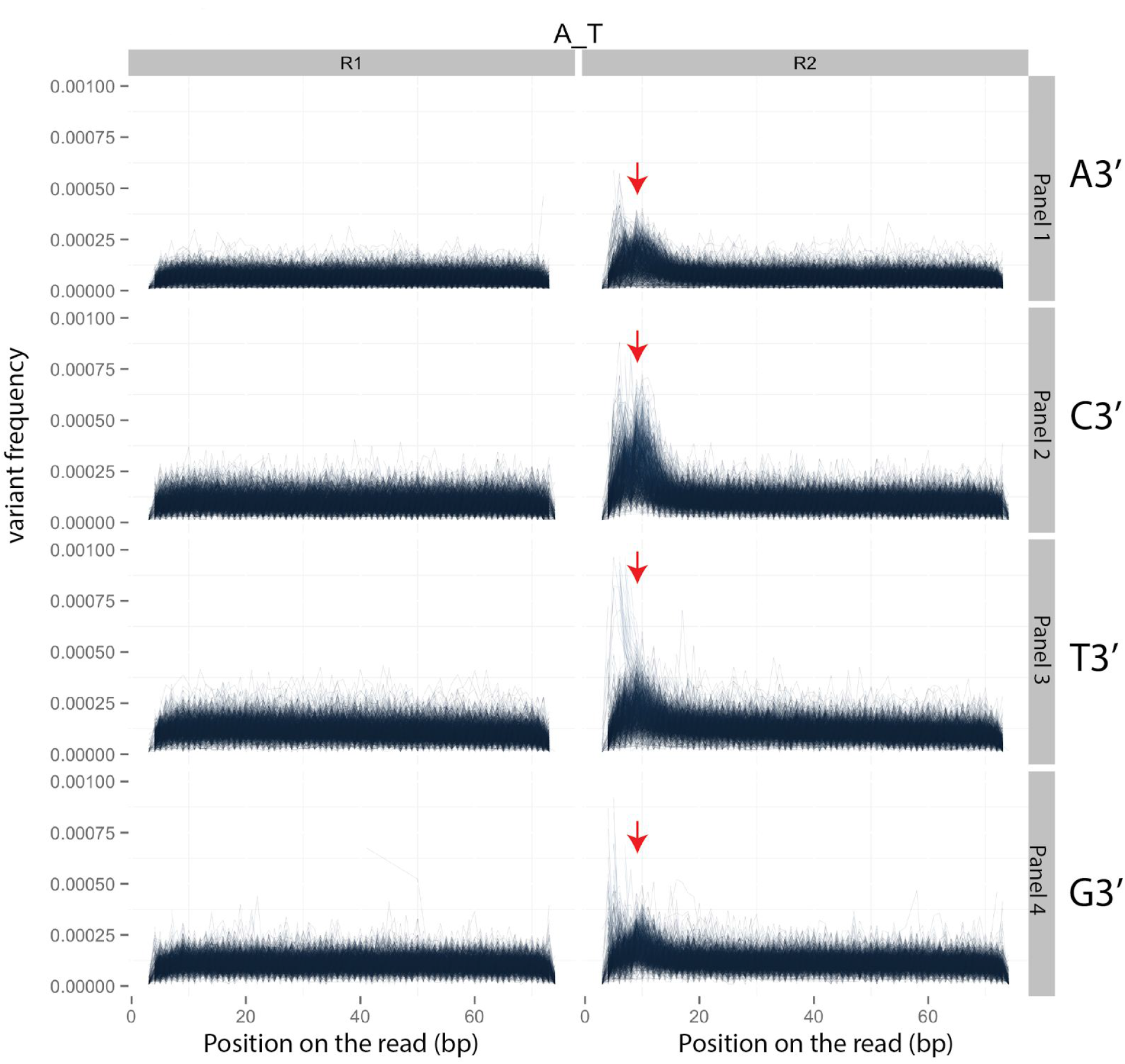
Damage in the TCGA dataset. A to T variant frequency as a function of read position in the R1 and R2 reads for sequence context ApA (panel1), ApC (panel2) and ApT (panel3) and ApG (panel4). Red arrows denote elevated variant frequency at the 5’ end of R2.

**Supplementary Figure 6.**
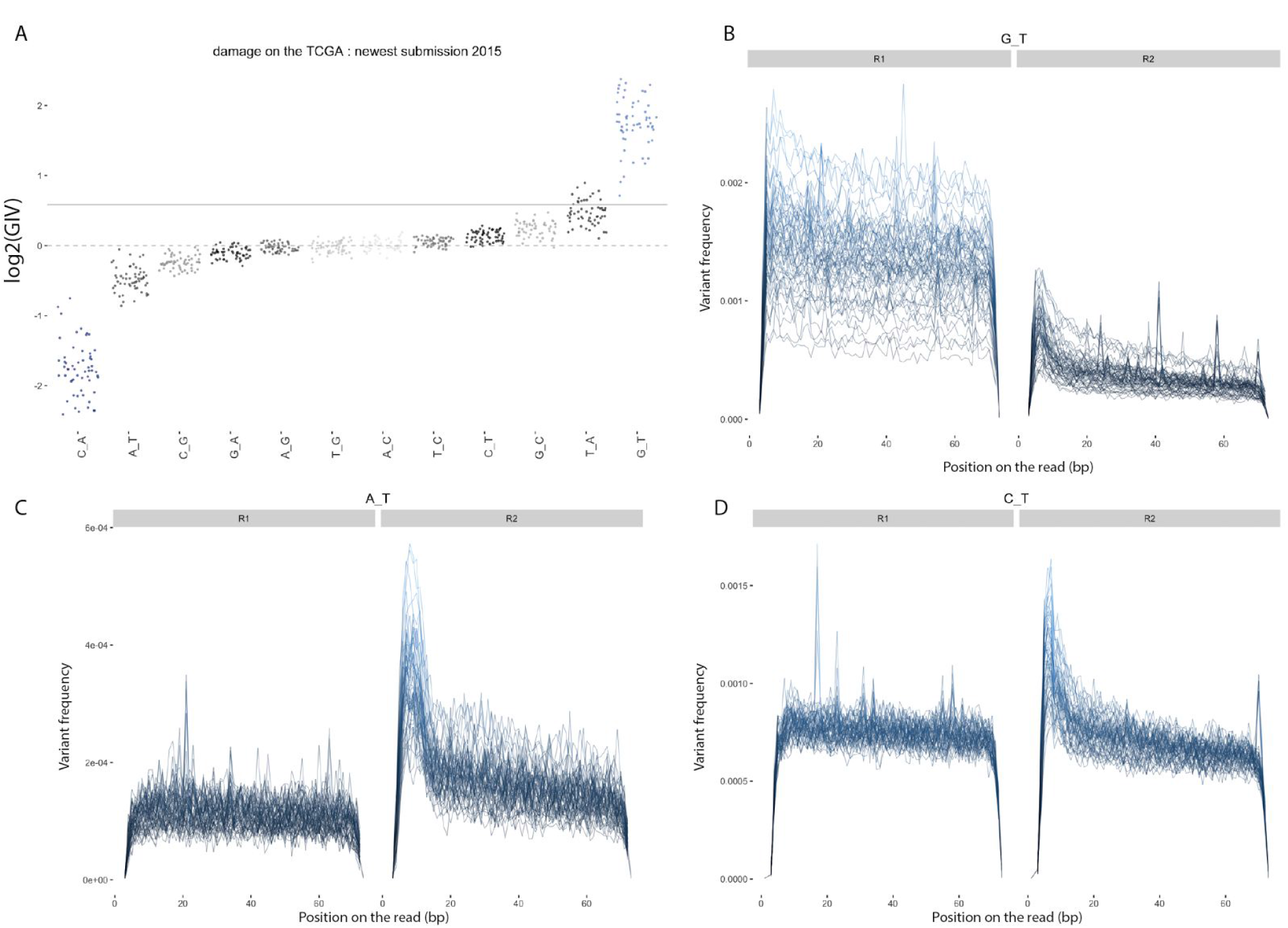
Damage in the most recently submitted TCGA runs. **A.** GIV scores for all the 12 nucleotide substitution classes. The dotted line denotes a GIV score of 1 which corresponds to no damage. The solid line denotes a GIV score of 1.5 corresponding to a limit of significant damage. **B.** G to T variation frequency as a function of the position on the read (bp) in R1 and R2 sequences. **C.** A to T variation frequency as a function of the position on the read (bp) in R1 and R2 sequences. **D.** C to T variation frequency as a function of the position on the read (bp) in R1 and R2 sequences.

**Supplementary Figure 7.**
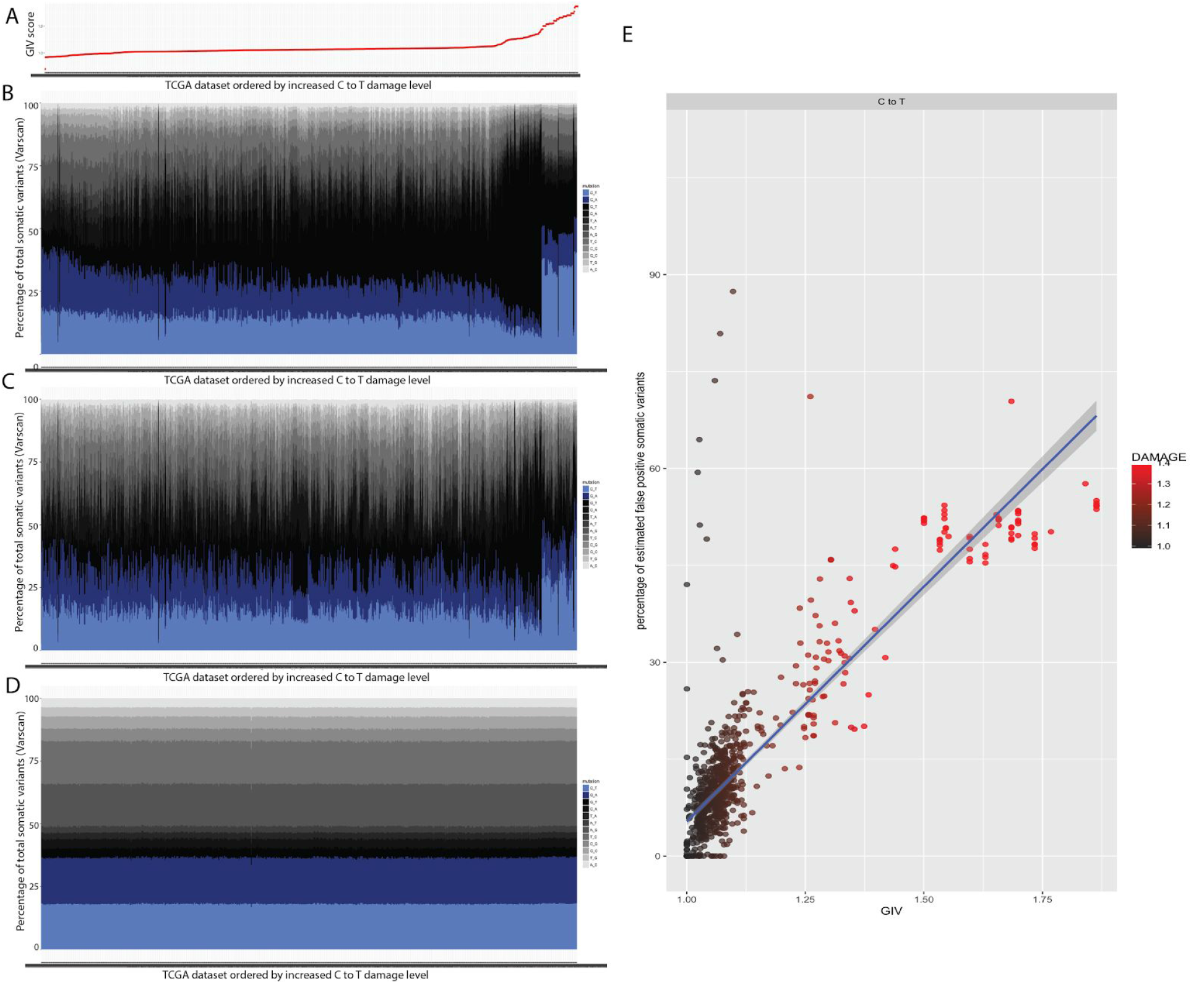
**A.** Distribution of the GIV_C_T_ scores in the ∼1800 tumor samples from the TCGA dataset. **B.** Profile of somatic variant calls for the ∼1800 tumor samples analysed and organized by increasing GIV_C_T_ score. The fraction of somatic variant calls of C to T types (light blue) is higher when compared to G to A (dark blue) for a subset of the datasets and the overall fraction of somatic variant calls of C to T types increase with the increase of GIV_G_T_ score. **C.** Same as in B except using the profile of high confidence somatic variant calls instead of somatic variant calls. **D.** Same as in B, except using the profile of germline variant calls instead of somatic variant calls. **E.** Estimated false positive rate of somatic variant calls of the C to T type (in percent) as a function of the GIV_C_T_.

**Supplementary table 1:** Shearing Buffer Composition

| Sample | Shearing Buffers |
| --- | --- |
| 1 | 1x TE (10 mM Tris pH8, 1 mM EDTA) |
| 2 | 10 mM Tris pH8, 0.1 mM EDTA |
| 3 | 10 mM Tris pH8 |
| 4 | 0.1x TE (1 mM Tris pH8, 0.1 mM EDTA) |
| 5 | 1 mM Tris pH8 |
| 6 | 1 mM EDTA pH8 |
| 7 | 0.1 mM EDTA pH8 |
| 8 | water |

In addition, we also detected several previously uncharacterized signatures of damage that correlate with buffering conditions during sonication. The first signature is characterized by an excess of A to T variants located on the first 20 bp of the 5’ end of the R2 sequences. This excess of A to T is reduced in repaired samples demonstrating that these variants are erroneous and result from DNA damage. Translated to the original DNA fragment, this damage leads to a T to A transversion at the 3’ end of the fragments. Similar to the G to T signature, this signature is stronger in an unbuffered shearing solution and diminishes with additional buffering capacity. Unlike G to T, this signature has strong context specificity with a 5’ T being the preferred context (*Supplementary figure 1B*). Other minor signatures include C to T and G to T variants located on the first 20 bp of the 5’ end of the R2 reads. Both signatures show strong context specificity (*Supplementary figure 1A and C*).

### Supplementary text 4. ClearSeq target enrichment and analysis

A panel of 151 cancer genes was captured using the ClearSeq Comprehensive Cancer panel (Agilent) and sequenced on an Illumina MiSeq platform using 2x75 paired-end protocol as described in Materials and Methods.

Trimming and mapping of the reads were done as described in the Materials and Methods section. The reads were locally remapped using GATK IndelRealigner. Reads were classified as R1 or R2 and variants were called using samtools mpileup with the following parameters: -l -O -s -q 10 -Q 30. We used the annotation file corresponding to the ClearSeq Comprehensive Cancer panel provided by Agilent for the -l parameter. For each genomic position, the coverage was downsampled to exactly 300 fold coverage by randomly sampling the mpileup file for all, R1 or R2 reads and the frequency of each variant was calculated.

### Supplementary Materials and Methods

#### Materials

All human genomic DNA in this study were obtained from BioChain Institute, Inc. (Newark, CA).

#### Illumina sequencing library preparation with and without DNA repair

All human genomic DNAs were sheared to 200 bp average fragment size using a Covaris S2 (Covaris Inc., Woburn, MA) with the following settings: 10% duty cycle, intensity 5, 200 cycles per burst and treatment time of 6 minutes. Without DNA repair, libraries were constructed using either the NEBNext DNA Library Prep Master Mix Set for Illumina or the NEBNext Ultra II DNA Library Prep Kit for Illumina (NEB, Inc.). Input DNA was 50 ng for all libraries unless otherwise mentioned. The fragmented DNA was blunted and dA tailed followed by adaptor ligation as described in the manufacturer’s manual. Library amplification was performed using 6-8 cycles of PCR using NEBNext Ultra II Q5 Master Mix. The library quality was assessed using a high sensitivity DNA chip on a Bioanalyzer (Agilent Technologies, Inc.). All libraries were indexed and paired-end sequenced on an Illumina MiSeq platform. For *in vitro* DNA repair, prior to the end repair step, the fragmented DNAs were treated by incubating the DNA with the repair mix at 200C for 15 minutes as described in the manual for the NEBNext FFPE DNA Repair Mix (NEB, Inc.). All other steps in the library preparation protocol were kept the same as for the unrepaired DNA library construction.

#### Deep sequencing of cancer gene panel from frozen human liver DNA

A panel of 151 cancer genes was captured from genomic DNA using the SureSelect Target Enrichment kit (Agilent) following the manufacturer’s protocol with modifications. DNAs were sheared in 0.1x TE buffer to 200bp using a Covaris S2 as described earlier. Briefly, 250 ng of fragmented genomic DNA was either treated with the NEBNext FFPE DNA Repair Mix followed by the NEBNext End Prep step of the NEBNext Ultra II DNA Library Prep workflow, or prepared without DNA repair using the same amount of DNA input and with the NEBNext Ultra II DNA Library Prep workflow. NEBNext adaptors and NEBNext multiplex PCR primers were used instead of the SureSelect adaptor mix and SureSelect ILM indexing pre-capture primers, respectively. Pre-captured libraries were amplified with 6 cycles of PCR using the NEBNext multiplex PCR primers and the NEBNext Ultra II Q5 Master Mix. Libraries were quantified on an Agilent BioAnalyzer and 750 ng was hybridized with an Agilent XT ClearSeq Comprehensive Cancer Panel. Post-captured DNA libraries were amplified with 14 PCR cycles using the same barcode index primers used prior to target enrichment. Two libraries (with and without DNA repair) were pooled and sequenced on an Illumina MiSeq sequencer at 2 x 75bp. Data from a total of four MiSeq runs were combined and analyzed for variant calling.

